# MEIS1 down-regulation by MYC mediates prostate cancer development through elevated HOXB13 expression and AR activity

**DOI:** 10.1101/848952

**Authors:** Nichelle C. Whitlock, Shana Y. Trostel, Scott Wilkinson, Nicholas T. Terrigino, S. Thomas Hennigan, Ross Lake, Nicole V. Carrabba, Rayann Atway, Elizabeth D. Walton, Berkley E. Gryder, Brian J. Capaldo, Huihui Ye, Adam G. Sowalsky

## Abstract

Localized prostate cancer develops very slowly in most men, with the androgen receptor (AR) and MYC transcription factors amongst the most well-characterized drivers of prostate tumorigenesis. Canonically, MYC up-regulation in luminal prostate cancer cells functions to oppose the terminally differentiating effects of AR. However, the effects of MYC up-regulation are pleiotropic and inconsistent with a poorly proliferative phenotype. Here we show that increased MYC expression and activity are associated with the down-regulation of *MEIS1*, a HOX-family transcription factor. Using RNA-seq to profile a series of human prostate cancer specimens laser capture microdissected on the basis of MYC immunohistochemistry, MYC activity and *MEIS1* expression were inversely correlated. Knockdown of *MYC* expression in prostate cancer cells increased expression of *MEIS1* and increased occupancy of MYC at the *MEIS1* locus. Finally, we show in laser capture microdissected human prostate cancer samples and the prostate TCGA cohort that *MEIS1* expression is inversely proportional to AR activity as well as *HOXB13*, a known interacting protein of both AR and MEIS1. Collectively, our data demonstrate that elevated MYC in a subset of primary prostate cancers functions in a negative role in regulating *MEIS1* expression, and that this down-regulation may contribute to MYC-driven development and progression.

## INTRODUCTION

Locally advanced prostate cancers harbor a limited number of recurrently altered genes whose expression change at the earliest stages of tumor development. These include down-regulation of the tumor suppressors *NKX3-1* and *PTEN* (often due to genomic deletion), up-regulation of *ERG* (due to fusion with *TMPRSS2*), and up-regulation of *MYC*, which often co-occurs with a single-copy gain of chromosome 8q24 [1–5]. Interestingly, up-regulation of *MYC* in most neoplastic tissues is a very early event that contributes to self-renewal and proliferation, but localized prostate cancer (PCa) is not hyperproliferative and focal amplification of *MYC* is rare [6, 7]. In part, the effects of the androgen receptor (AR) in terminally-differentiated luminal prostate cells are disrupted by MYC and other cofactors including FOXA1 and HOXB13 to re-engage proliferative processes during tumorigenesis [7–11].

Recently, increased awareness that the vast majority of prostate cancers are indolent has led to increased molecular profiling of tumor biopsies prior to definitive treatment. Although MYC expression has been observed in the cancer precursor high-grade prostatic intraepithelial neoplasia, MYC expression in indolent-appearing tumor cells predicts the presence of higher-grade disease and is associated with poor differentiation [3–5]. In localized prostate cancers, up-regulated MYC has been further associated with alterations in nucleoli structure, concomitant with increased biogenesis of ribosomal RNA, increased purine metabolism and expression of the telomere RNA subunit *TERC* [12–14]. These effects contrast sharply with the phenotype of highly amplified MYC, which is enriched in metastatic and treatment-resistant prostate cancers, and mirrors the role of MYC in other cancer types via effects on AKT to contribute towards cell division and survival [15–17]. Nonetheless, even in primary PCa, MYC protein expression is diffuse and heterogeneous [3].

We sought to examine the genetic contribution of MYC to the development and progression of primary PCa in the context of dysregulated growth by using anti-MYC immunohistochemistry and performing laser capture microdissection on populations of human prostate tumor cells with varying expression of MYC protein. We show using transcriptome profiling that increased MYC activity is inversely correlated with expression of Myeloid Ecotropic viral Insertion Site 1 (*MEIS1*), a transcription factor that interacts with and regulates the activity of HOX homeodomain transcription factors, including HOXB13 [18–20]. We determined that MYC binding to the *MEIS1* locus decreases as MYC levels increases, and that *MEIS1* expression is negatively correlated with *HOXB13* expression and AR activity.

## METHODS AND MATERIALS

### Study approval

This research was conducted in accordance with the principles of the Declaration of Helsinki. All patients provided informed consent prior to participating tissue procurement protocols, and all samples were deidentified as per institutional policies. The collection of radical prostatectomy specimens was approved by the Dana Farber/Harvard Cancer Center Institutional Review Boards, under protocol numbers 11-250, 15-008, and 15-492. The collection of prostate biopsy specimens was approved by the National Institutes of Health Institutional Review Board, under protocol number 15-c-0124. Tissues were fixed in formalin and embedded in paraffin according to standard procedures.

### Pan-cancer analysis

Gene-level normalized FPKM expression values for The Cancer Genome Atlas (TCGA) RNA-seq data were downloaded from the NCI Genomic Data Commons (https://gdc.cancer.gov). Cases were filtered for tumor samples within each organ type and cases with missing gene expression values were removed. Single-sample gene set enrichment analysis was performed on the GenePattern server (https://cloud.genepattern.org) using the ssGSEAProjection module version 9.1.1 with the following parameters: weighting exponent: 0.75; combine mode: combine.add; sample normalization method: none. For each TCGA tumor type, ssGSEA projection values were obtained for a 17-gene MYC-independent proliferation signature and a 54-gene MYC activity signature [21].

### Immunohistochemistry (IHC)

Serial sections of formalin-fixed, paraffin-embedded tissues were cut at 6 μm thickness onto Superfrost Plus slides (Fisher Scientific). Slides were stained with the following: hematoxylin & eosin (H&E); anti-ERG (EPR3864; Abcam Cat# ab92513, 1:100 dilution), anti-MYC (Y69; Abcam Cat# ab32072, 1:100 dilution); anti-Ki67 (D2H10; Cell Signaling Cat# 9027, 1:100 dilution), anti-PTEN (D4.3; Cell Signaling Cat#9188, 1:100 dilution); and PIN-4 cocktail (ready-to-use; Biocare Medical Cat# PPM225DS). Antigen retrieval was performed using a pre-heated steamer for 20 minutes in Diva Decloaker (Biocare Medical, Cat# DV2004MX) for Ki67, ERG, PTEN and PIN-4 or High pH Antigen Retrieving Solution (Abcam, Cat# 972) for MYC, followed by 20 minutes of cooling. Sections were blocked with hydrogen peroxide (Sigma-Aldrich, Cat# 216763) for 5 minutes, blocked with Background Sniper (Biocare Medical, Cat# BS966) for 10 minutes for PIN-4 or VectaStain Elite ABC HRP kit (Vector Laboratories, Cat# PK-6101) for MYC, Ki67, PTEN and ERG, and incubated with primary antibody overnight at 4°C (MYC, Ki67, ERG and PTEN) or for one hour (PIN-4). Secondary labeling was performed using Mach 2 Double Stain for PIN-4 (Biocare Medical, Cat# MRCT525) or the VectaStain Elite ABC HRP kit for MYC, Ki67, ERG and PTEN for 30 minutes. Avidin-biotin complexing was then performed for 30 minutes for MYC, Ki67, ERG and PTEN. Colorimetric detection was achieved using DAB Peroxidase HRP (Vector Laboratories, Cat# SK4100) for MYC, Ki67, ERG and PTEN, Betazoid DAB (Biocare Medical, Cat# BDB2004) for PIN-4, and Vulcan Red Fast Chromogen (Biocare Medical, Cat# FR805) for PIN-4. Counterstaining was performed using Mayer’s Hematoxylin Solution (Sigma Aldrich, Cat# MHS16). PIN-4 stained slides were air-dried. MYC, Ki67, ERG and PTEN stained slides were dehydrated through graded alcohol and cleared in xylenes. Slides were mounted using Permount (ThermoFisher).

### Computer-aided image analysis

Stained slides were scanned using the 20× objective (Plan-Apochromat, NA 0.8) with brightfield illumination on an AxioScan.Z1 (Zeiss). All slides were reviewed by a board-certified surgical pathologist following the 2014 ISUP guidelines [22]. Whole slide. CZI files of Ki67- and MYC-stained slides were imported into Definiens Developer XD 64. The magnification for each analysis was set to 40×, with 0.11 μm/pixel for both Ki67 and MYC solutions. IHC stain was identified as brown chromogen. The tumor cells were counted using their nuclear stain.

For Ki67, composer magnification was set to 6× with 12 training subsets at segmentation level 9. Segments were classified as tumor or stroma, with normal glands excluded. Within the tumor pattern, cellular analysis magnification was set to 10× with 12 training subsets, and within nuclear detection, thresholds were 0.2 for hematoxylin and 0.5 for brown chromogen with typical nuclear size set to 30 μm. Nucleus classification: low vs. medium at 0.65; medium vs. high at 0.86.

For MYC analysis, composer magnification was set to 7.5× with 12 training subsets at segmentation level 7. Segments were classified as tumor or stroma, with normal glands excluded. Within the tumor pattern, cellular analysis magnification was set to 10× with 12 training subsets, and within nuclear detection, thresholds were 0.1 for hematoxylin and 0.45 for brown chromogen: 0.45 with typical nuclear size set to 30 μm. Nucleus classification: low vs. medium at 0.5; medium vs. high at 0.7.

The total number of positively stained nuclei were reported, along with distribution of low-, medium-, and high-intensity stained nuclei for each tumor focus. A percent positive index score was calculated using a weighted average divided by the total number of nuclei, where *index* = [ (1 × nuclei stained low) + (2 × nuclei stained medium) + (3 × nuclei stained high)] ÷ (3 × total nuclei).

### Cell lines and derivatives

RNA knockdown of *MYC* expression was achieved using the SMARTvector lentiviral shRNA system (Dharmacon) and a standard second-generation packaging system. hCMV-TurboGFP targeting vectors were MYC 512: V3SH11240-225190433 (GGTCGATGCACTCTGAGGC, targeting ORF); MYC 627: V3SH11240-226339523 (TTGATCATGCATTTGAAAC, targeting 3’ UTR); MYC 637: V3SH11240-226439183 (GTAGAAATACGGCTGCACC, targeting ORF); and non-targeting control: VSC11707. Briefly, 293FT cells (ThermoFisher) were reverse-transfected with a lentiviral shRNA expression vector in a 10 cm dish, using envelope (pCMV-VSV-G, #8454, Addgene) and packaging (pCMV-dR8.2 dvpr, #8455, Addgene) vectors following the ThermoFisher ViraPower protocols. Media containing lentivirus were harvested two days post-transfection, passed through a 0.45 μm filter, aliquoted and frozen. HT-1080 cells (ATCC) were used for titering virus following the ThermoFisher ViraPower protocol.

LNCaP cells (clone FGC, catalog number CRL1740) were purchased from ATCC. Cell line authentication by STR profiling and mycoplasma testing was performed every six months (Laragen, Inc). LNCaP cells were seeded in 6-well plates at 1×10^5^ cells per well and transduced with lentivirus at a multiplicity of infection of 1 in complete media for 24 hours. Complete media were then replaced one day post-transduction, and selection with puromycin (1 μg/mL, Life Technologies) began two days post-transduction. Transduction was confirmed by visual confirmation of GFP fluorescence in >90% of cells, and knockdown was confirmed by qPCR and Western blotting.

### Laser capture microdissection and RNA purification

Laser capture microdissection (LCM) was performed as previously described [23]. For within-patient analyses (high and low MYC levels in the same patient), cases were selected based on having regions of differential MYC staining intensity by IHC, but concordant ERG and PTEN status by IHC. Areas of high MYC were defined by moderate (++) or high (+++) staining in at least 30% of cancer cells, while areas of low MYC were defined by weak (+) or absent (-) staining in at least 75% of cancer cells, with intense staining limited to 10% of cancer cells. LCM was guided by review of serially-sectioned slides stained with MYC, visualized using ZEN Browser (Zeiss) on an adjacent monitor as reference. High or low MYC cancer glands from each case were collected on separate caps and a photomicrograph was acquired for the purposes of estimating tumor cell purity in each sample. For between-patient analyses, MYC status was ascertained by IHC and assigned to the foci of tumor cells subjected to LCM. The distinction between within-patient and between-patient analyses is that all within-patient cases harbored both MYC-high and MYC-low populations of tumor cells.

RNA was extracted using the RNeasy FFPE Kit (Qiagen) following the manufacturer’s protocol with modifications. Briefly, the film to which LCM cells adhered was removed using a disposable blade, immersed into Buffer PKD with proteinase K, digested for 15 min at 56°C, followed by a 15-minute incubation at 80°C. After centrifugation and DNase treatment to remove genomic DNA, concentrated RNA was purified using RNeasy MinElute spin columns and eluted with 30 μL RNase-free water. RNA yields were quantified using RiboGreen reagent (Life Technologies).

### Quantitative polymerase chain reaction

RNA was purified using the RNeasy Plus Mini Kit (Qiagen) and quality tested using the High Sensitivity RNA ScreenTape (TapeStation 4200, Agilent Technologies). Expression levels of *MYC* and *MEIS1* transcript were quantified using TaqMan Fast Virus 1-Step Master Mix (Life Technologies) using PrimeTime qPCR Probe Assays from Integrated DNA Technologies in a 3:1 ratio of primer to 5′ 6-FAM and 3′ TAMRA labeled probe. Sequences of the primer-probe sets are listed in Supplementary Table 1. qPCR was performed in duplex with GAPDH endogenous control TaqMan assays (Life Technologies), and relative transcript levels were quantified using the 2^−ΔΔCT^ method [24].

### Western Blotting

Total cell lysates were isolated using RIPA buffer (Pierce) containing protease and phosphatase inhibitors (Pierce). 15 μg of soluble protein per well were separated by SDS-PAGE and transferred to nitrocellulose membranes. Blots were blocked for 1 hour in 5% nonfat dry milk in TBS/0.05% Tween-20 (TBS-T) and incubated with primary antibodies at 4°C overnight. Primary antibodies used were anti-MYC (Y69; ab32072, Abcam, 1:2000 dilution) and anti-GAPDH (6C5; MAB374, Millipore Sigma, 1:1000 dilution). After washing with TBS-T, the blots were incubated with anti-rabbit or anti-mouse horseradish peroxidase-conjugated secondary antibody (Jackson ImmunoResearch, 1:10,000 dilution) for 1 hour and washed 4 times. Chemiluminescence was detected with Western Lightning Plus ECL (Perkin Elmer) and recorded using a ChemiDoc Imaging System (Bio-Rad).

### ChIP-seq Analysis

ChIP was performed according to the ChIP-IT High Sensitivity Kit (Active Motif) protocol. Briefly, cultured LNCaP cells with control or MYC knockdown constructs were crosslinked, quenched, lysed, and the chromatin sheared to a size between 200-1000 bp. *Drosophila melanogaster* chromatin and antibody spike-in controls were added per the protocol. For each ChIP, 30 μg of chromatin were incubated with anti-MYC antibody (ab32072, Abcam, 1:100 dilution) on an end-to-end rotator overnight at 4°C. Antibody-bound protein/chromatin complexes were immunoprecipitated with Protein A/G beads, reverse-crosslinked, and DNA purified. Quality of ChIP-enriched DNA was assessed using the ChIP-IT qPCR Analysis Kit (Active Motif).

Purified ChIP DNA was assembled into libraries using the TruSeq ChIP Library Preparation Kit (Illumina) according to the manufacturer’s protocol, pooled in equimolar ratios, and sequenced on a NextSeq 500 (Illumina) with 76 cycles of single-end sequencing. All samples had 90% of bases at Q30 or above, with yields between 26-42 million pass filter reads per sample. Samples were adaptor-trimmed with Trim Galore (https://github.com/FelixKrueger/TrimGalore) and reads were mapped to the human genome GRCh37 and *Drosophila melanogaster* genome dm3 with BWA-MEM [25]. Aligned reads were deduplicated using Picard Tools (https://broadinstitute.github.io/picard). Peak calling was performed using MACS2 [26] callpeak with the parameters -g hs and -q 0.01, differential binding of peaks was ascertained using DiffBind [27] with default parameters, and motif analysis was performed using HOMER [28] findMotifsGenome.pl with the parameters -size 200 and -len 8,10,12. To normalize for technical and genome-wide variability, reads mapping to dm3 were used to calculate reads per by million mapped dm3 reads (RRPM) as described [29].

### Gene expression analysis

Whole transcriptome profiling of LCM tissue was performed at the CCR Illumina Sequencing Facility using the Illumina HiSeq v4 sequencing system to a depth of 50 million reads (50 cycles, paired-end). Sample reads were trimmed for adapters using Trim Galore before alignment to the GRCh37 reference genome using STAR [30] with default parameters. Aligned reads were quantified using featureCounts [31] with default parameters, and differential expression analysis between MYC-high and MYC-low samples was performed using edgeR [32]. The log_2_ counts per million (CPM) values from the output were used for downstream analysis.

From the TCGA PRAD dataset used in the pan-cancer analysis, the 20 cases with highest MYC activity signature and the 20 cases with the lowest MYC activity signature were re-analyzed using the same pipeline as tissue for determining differentially expressed genes. FASTQ files for those cases were retrieved from the NCI Genomic Data Commons via access to dbGaP phs000178.

Gene set enrichment analysis (GSEA) was performed in clusterProfiler [33] comparing MYC-high versus MYC-low LCM foci and TCGA cases, run against gene sets in the Molecular Signature Database (v6.2).

### Statistical Analysis

Statistical analyses were performed using GraphPad Prism 8 for Mac. Statistical tests used and relevant variables are indicated in the legend of each figure.

## RESULTS

### MYC activity is weakly associated with proliferation in primary prostate cancer

Although MYC orchestrates a broad range of biological functions, it has been shown in many cancers that MYC drives tumorigenesis by potentiating or stimulating cell growth and proliferation [34]. To assess the relationship between MYC activity and proliferation across cancer types, we evaluated MYC activity and cell proliferation rate in The Cancer Genome Atlas (TCGA) pan cancer cohort using single-sample gene set enrichment analysis (ssGSEA) with MYC activity and proliferation signatures [21]. As shown in Figure 1A, signatures for MYC activity and proliferation were correlated in all cancers with the vast majority showing strong correlation (*r* > 0.5). However, prostate adenocarcinoma (PRAD) was amongst the lowest correlating cancer types (*r* = 0.4822, 95% C.I. 0.4112 to 0.5472; *P* < 0.0001). To validate this observation, we screened a new cohort of treatment naïve Gleason score 7 prostate tumors for MYC and Ki67 expression by immunohistochemistry (IHC). Although MYC was expressed in a majority of tumor cells, Ki67 expression was limited to < 1% of cells (Fig. 1B, left). We applied automated image analysis to quantify millions of nuclei across 20 tumor foci from 10 different cases. While the MYC IHC index ranged from 0 (low expression) to 0.46 (high expression), the Ki67 index was less than 0.1 in 18 out of 20 foci (Fig. 1B, right). An unweighted Pearson correlation demonstrated that MYC IHC was weakly but positively correlated with Ki67 (*r* = 0.4954, 95% C.I. 0.06772 to 0.7693, P = 0.0263), similar to what we observed in TCGA, indicating that these transcriptional signatures accurately recapitulated MYC and Ki67 protein abundance. Because high Ki67 expression was not associated with high MYC expression, proliferation may have been dependent upon MYC activity, but MYC expression alone was not sufficient for proliferation.

**Figure 1.**
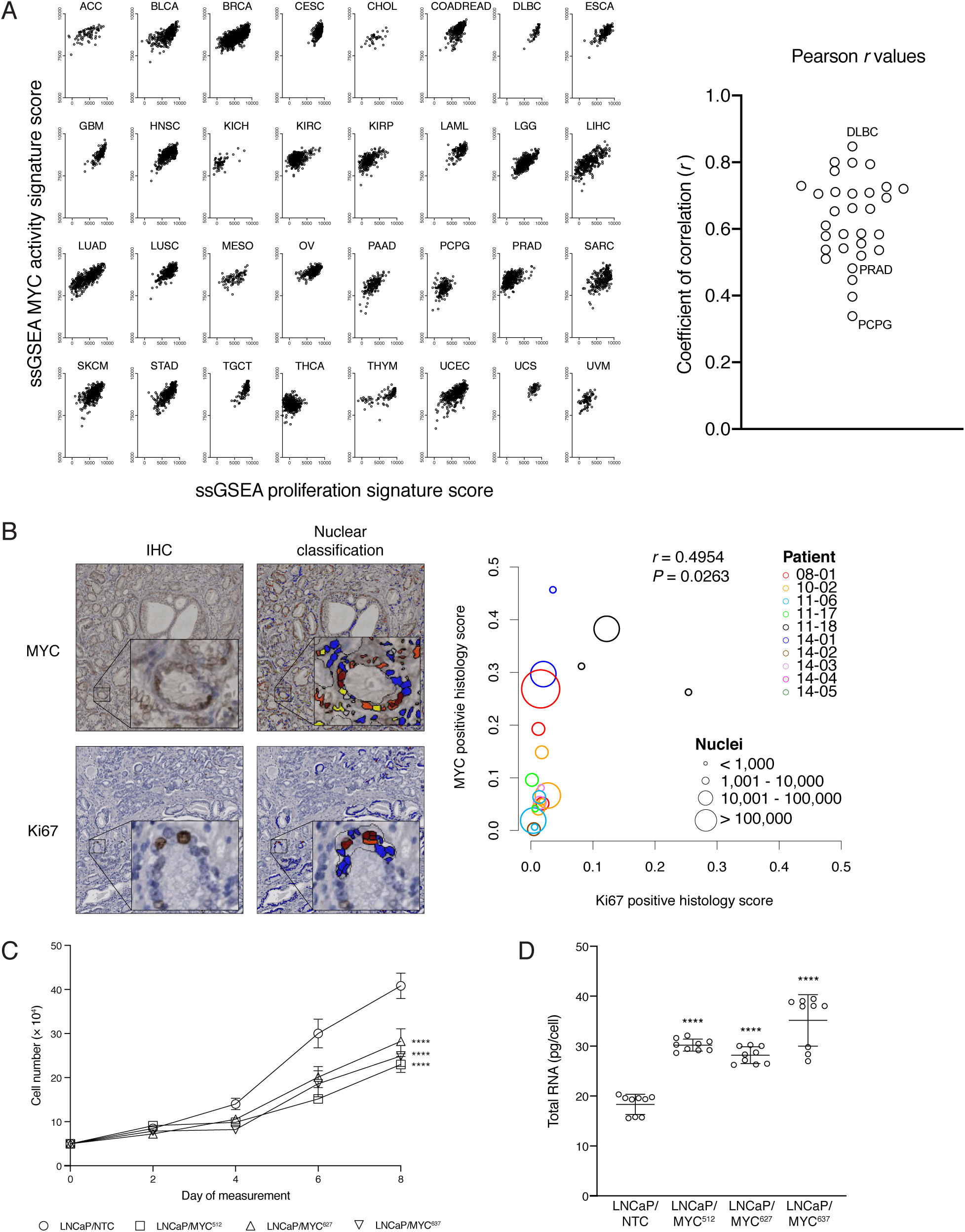
MYC and proliferation in cancers. A. Left: correlation of the 54-gene ssGSEA MYC activity signature score and the 17-gene ssGSEA proliferation signature score for each tumor type from the TCGA pan-cancer cohort. All scatter plots are on the same scale. Right: plot of Pearson *r* coefficients of correlation from each scatter plot shown on the left. All tumor types showed a positive correlation, ranging from 0.3386 (PCPG: pheochromocytoma and paraganglioma) to 0.8473 (DLBC: diffuse large B-cell lymphoma). PRAD: prostate adenocarcinoma, *r* = 0.4822. All correlations were significant at *P* < 0.001 except THCA (thyroid carcinoma; *P* = 0.5543). B. Left: representative IHC staining used as input for automated image analysis with Definiens Tissue Studio and the processed file used for quantification of histologic score via nuclear classification, for anti-MYC and anti-Ki67 immunostains. Right: unweighted scatter plot and Pearson correlation of the quantified intensities of anti-Ki67 versus anti-MYC stain. Each open circle represents a contiguous quantified region, color-coded by patient. Multiple regions from the same patient indicate multiple noncontiguous regions were measured, and the size of each circle indicates the number of nuclei considered for the analysis. C. Assessment of cell growth *in vitro* as a function of *MYC* knockdown. LNCaP cells expressing non-targeting or MYC-targeting hairpins were seeded at a density of 50,000 cells per well and counted every 2 days in triplicate using a hemocytometer by three different individuals blinded to the identities of the cultures. Data shown represents the mean ± standard error of ten independent experiments. **** *P* < 0.0001 from control. D. RNA content as a function of *MYC* knockdown. Total RNA was extracted from 1 × 10^6^ pelleted LNCaP cells expressing non-targeting or *MYC*-targeting hairpins. RNA concentrations were measured by Nanodrop and total RNA was calculated per cell. Data represents mean ± standard deviation of 3 independent experiments conducted in triplicate, with open circles representing individual replicate RNA quantity values (****, *P* < 0.0001).

To further examine the role of MYC in prostate tumorigenesis, we knocked down MYC in LNCaP cells which had physiologically-elevated levels of MYC. Like primary PCa, LNCaP cells express AR, are androgen sensitive, and grow slowly despite elevated MYC [35]. Using MYC-knockdown cells as a model of MYC-low prostate cancer cells, we show in Supplementary Figure 1 that we achieved ~50% reduction in *MYC* transcript and MYC protein expression. As anticipated, knockdown of *MYC* was associated with longer intervals between cell division *in vitro* (Fig. 1D) but the decrease in cell growth was not proportional to the decrease in *MYC* expression. The relationship between MYC and proliferation in other cancer types also stipulates a role for MYC as a universal amplifier of transcription, [36] alleviating constraints on cell growth and proliferation [37]. However, we observed the opposite: the total amount of cellular RNA increased when *MYC* was knocked down in LNCaP cells (Fig. 1D). Together, these findings indicate that the cellular behavior of MYC in prostate cancer contrasts with other tumor types, and that MYC does not act solely in a proliferative capacity.

### Increased MYC is associated with specific up- and down-regulation of target genes

We hypothesized that analysis of tumors with different levels of MYC expression would identify genes that may contribute to MYC activity in PCa pathogenesis. Using samples of human radical prostatectomy specimens stained with anti-MYC, we identified concomitant regions of high and low MYC expression in the same patients. We subjected these within-patient sets of tumor foci to laser capture microdissection (Fig. 2A), controlling for PTEN and ERG status by confirming their concordance within each case. Of the 19 foci microdissected, we designated 42% MYC-high (*n* = 8) and 58% MYC-low (*n* = 11). We performed RNA-seq on these cases and derived a limited gene set of 303 up- or down-regulated genes, with only 17 genes showing expression changes of 4-fold or more with adjusted *P* values less than 0.1 (Fig. 2B). Consistent with our observation of increased mRNA in LNCaP cells expressing MYC knockdown hairpins (see Fig. 1D), we observed more genes downregulated than upregulated in MYC-high vs. MYC-low tumor foci.

**Figure 2.**
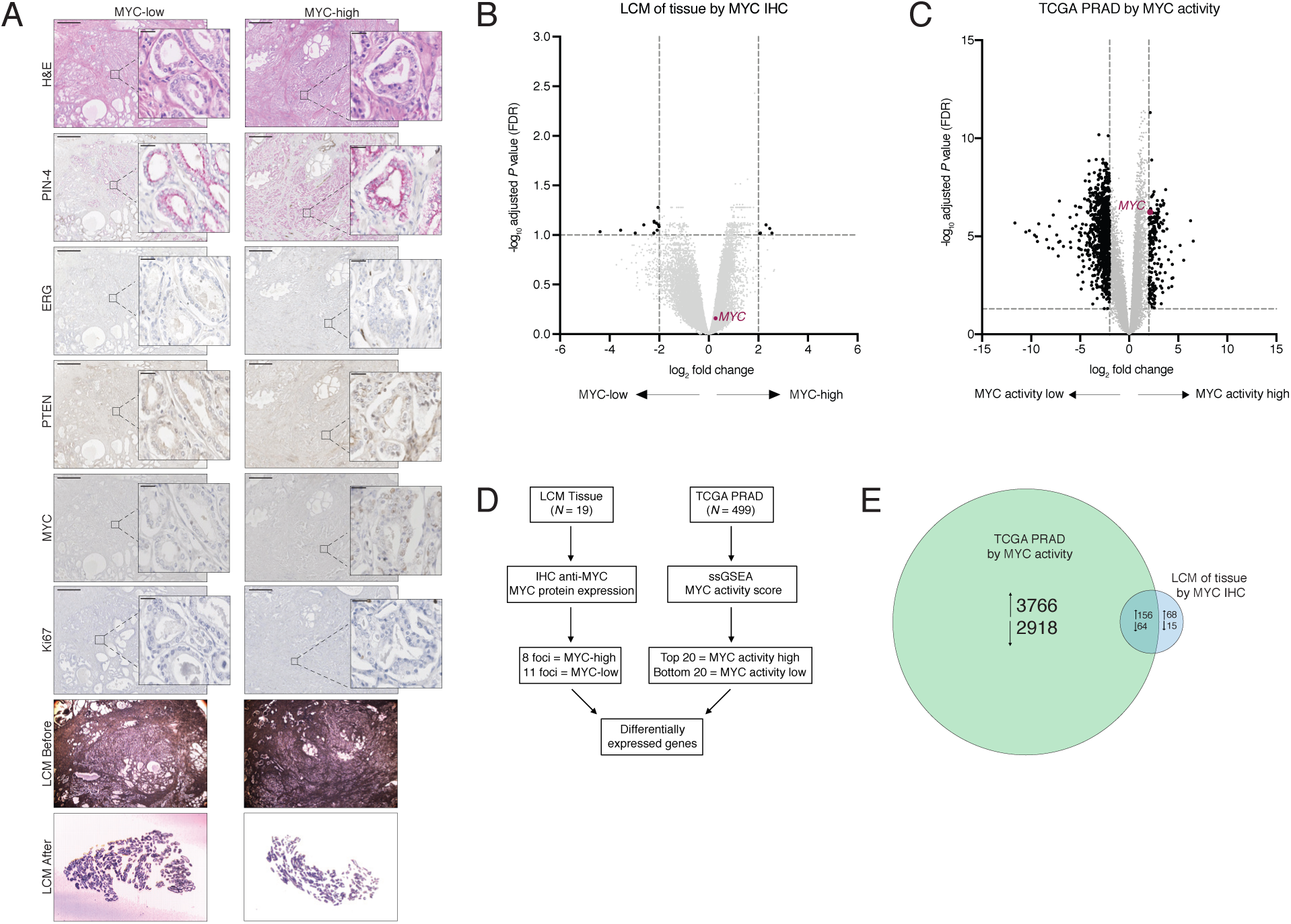
Effect of MYC levels on gene expression. A. Representative IHC of case 14-03 showing heterogeneity of MYC expression within a single patient. Serial sections were stained with H&E and PIN-4 cocktail to distinguish benign glands and high grade prostatic intraepithelial neoplasia from tumor foci; anti-ERG and anti-PTEN immunostains were performed to select regions with concordant phenotypes. Micrographs of stained tissue before and after laser capture microdissection (LCM) are also shown. Scale bar: 500 μm; inset scale bar: 20 μm. B. Volcano plot depicting differentially expressed genes in MYC-high vs. MYC-low tumors in the LCM cohort, with the false discovery rate (adjusted −log_10_ *P* value) on the *y*-axis and the log_2_-transformed fold-change (MYC-high vs. MYC-low) on the *x*-axis. Expression of *MYC* is shown in red. Black dots represent genes with fold change ≥ ±4 and adjusted *P* value < 0.1. C. Volcano plot depicting differentially expressed genes in MYC activity high vs. MYC activity low tumors in the TCGA PRAD cohort, with the false discovery rate (adjusted −log_10_ *P* value) on the *y*-axis and the log_2_-transformed fold-change (MYC activity high vs. MYC activity low) on the *x*-axis. Expression of *MYC* is shown in red. Black dots represent genes with fold change ≥ ±4 and adjusted *P* value < 0.05. D. Flowchart of the strategy used to identify lists of differentially expressed genes from the LCM and TCGA PRAD cohorts. E. Euler diagram depicting distinct and shared differentially expressed genes between the LCM and TCGA PRAD cohort. Circles are drawn to scale.

To further refine genes of potential interest, we analyzed a second dataset comprised of whole transcriptome sequencing from 499 primary tumors in the TCGA PRAD cohort. Using the mRNA expression values of genes in the MYC activity signature (see Fig. 1A), we compared the top 20 and bottom 20 cases based on average median absolute deviation-modified *z*-score (Supplementary Fig. 2), and established a second list of differentially expressed genes (Fig. 2C). As these cases were compared between patients rather than within-patient, a far greater number of genes were differentially expressed (1064) with fold-change of at least 4 and adjusted *P* values less than 0.05.

When comparing differentially expressed genes between MYC-high vs. MYC-low tumors, *MYC* expression was consistently greater in the MYC-high group in the LCM primary PCa cohort (Fig. 2C and Supplementary Table 2). As MYC RNA and protein expression often do not correlate due to the tight posttranslational regulation of MYC [38, 39], the statistically insignificant increase in MYC expression in the MYC-high group was expected. However, MYC expression was amongst the upregulated genes in the MYC activity high group, confirming proper stratification in the TCGA PRAD cohort as well (Fig. 2D and Supplementary Table 3). Without any lower-bound cut-off on expression fold-change, we identified 305 and 6,907 significant differentially expressed genes between MYC-high and MYC-low prostate tumors in the LCM tissue and TCGA PRAD cohorts, respectively (Fig. 2E). Of these, only 220 were in common at a false discovery rate threshold of 0.1 for LCM tissue and 0.05 for TCGA PRAD (Fig. 2E, and Supplementary Table 4).

### MEIS1 is negatively associated with MYC expression and MYC activity

Given that we observed many more genes downregulated in MYC-high tumors than we would have expected if MYC were functioning as a genome-wide transcriptional amplifier, we hypothesized that collective analyses of gene expression might reveal coordinated down-regulation of biological processes that may contribute to MYC-driven PCa tumorigenesis. We therefore performed gene set enrichment analysis (GSEA) using both the LCM and TCGA cohort datasets as inputs, limiting our analyses to gene sets containing at least one differentially expressed gene shared by the TCGA and LCM cohorts. As depicted in Figure 3A, our initial comparative analyses demonstrated significant and concordant enrichment of 1737 gene sets (Supplementary Table 5). To narrow our focus further, we refined our search only to include gene sets that incorporated at least one target of master regulators on the premise that increased MYC activity would misregulate a multitude of pathways. These analyses identified 10 gene sets, a preponderance of which were associated with development and survival (Table 1).

**Table 1.** List of common gene sets negatively enriched in MYC-activity high vs. low comparing LCM tissue and TCGA cohorts. NES: normalized enrichment score.

| mSigDB Gene set | LCM Tissue |  | TCGA |  |
| --- | --- | --- | --- | --- |
|  | *NES* | *Q value* | *NES* | *Q value* |
| Takeda Targets of NUP98 HOXA9 Fusion 10D DN | -2.079 | 0.014 | -2.004 | 0.005 |
| Hecker IFNB1 Targets | -2.103 | 0.014 | -2.096 | 0.005 |
| Tonks Targets of RUNX1 RUNX1T1 Fusion HSC DN | -2.117 | 0.014 | -2.030 | 0.005 |
| Acevedo FGFR1 Targets in Prostate Cancer Model DN | -2.136 | 0.014 | -2.275 | 0.005 |
| Kim GLIS2 Targets UP | -2.189 | 0.014 | -2.113 | 0.005 |
| Servitja Iselt HNF1A Targets UP | -2.266 | 0.014 | -2.102 | 0.005 |
| Vilimas NOTCH1 Targets UP | -2.310 | 0.014 | -2.237 | 0.005 |
| Phong TNF Targets UP | -2.349 | 0.014 | -2.180 | 0.005 |
| Onder CDH1 Targets 3 DN | -2.437 | 0.014 | -2.181 | 0.005 |
| Gaurnier PSMD4 Targets | -2.608 | 0.014 | -2.307 | 0.005 |

**Figure 3.**
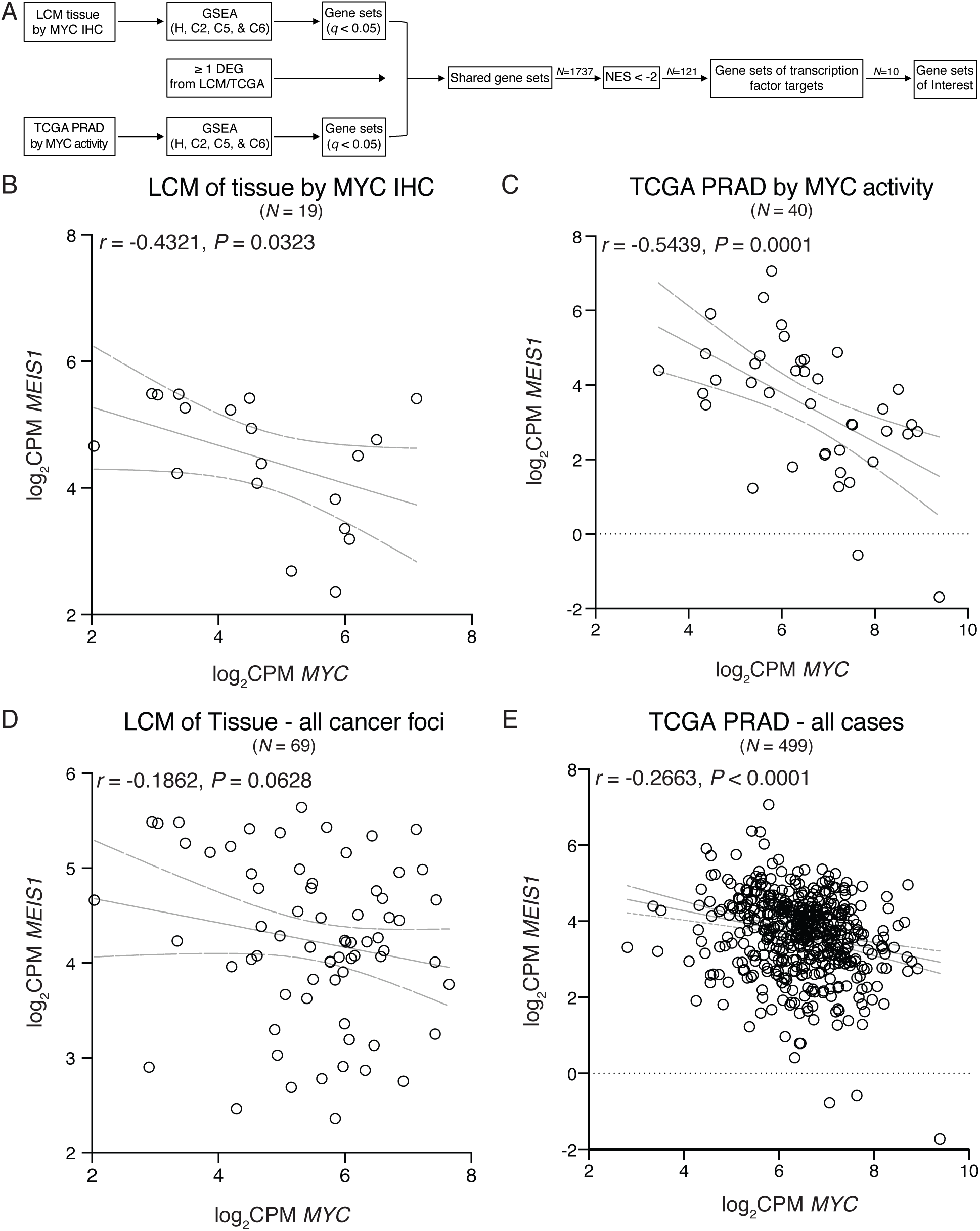
Association between MYC and MEIS1 expression. A. Flowchart of the strategy used to identify potential MYC target pathways based on differentially expressed genes. B-E. Correlation of the log_2_ CPM expression levels between *MYC* and *MEIS1* in the initial LCM cohort (*N* = 19, B), the initial TCGA PRAD cohort (*N = 40*, C), the entire LCM primary PCa cohort (*N* = 69, D), and the entire TCGA PRAD cohort (*N* = 499, E). The gray dotted lines represent the 95% confidence intervals for each correlation.

Of note, gene sets related to NOTCH, RUNX1, HOXA9, and TNF were negatively enriched in MYC-high vs. MYC-low tumors. This suggested that one or more transcription factors exert effects in opposition to MYC, and may by guiding MYC-mediated misregulation of these pathways. In particular, the transcription factor MEIS1 was identified as a common and important regulator in each of the aforementioned pathways. For example, MEIS1 has been shown to regulate genes in the NOTCH pathway [40] and sensitizes cells to TNF [41]. Moreover, MEIS1 is essential for the expression of genes driven by the HOXA9-NUP98 fusion in acute myeloid leukemia [42–44]. Therefore, we examined whether *MEIS1* expression was associated with increased MYC activity as a proxy for its role in PCa development.

We observed that *MEIS1* expression was reduced in MYC-high cases compared to MYC-low tumors, for both the LCM and TCGA cohorts (Supplementary Tables 2 and 3, respectively). At a case-by-case level, *MEIS1* expression was negatively correlated with *MYC* (Fig. 3B) and MYC activity (Fig. 3C) in the LCM and TCGA cohorts, respectively. To assess whether this association occurred in unselected populations of primary PCa, we analyzed RNA-seq data from an additional 69 microdissected tumor foci and the entire TCGA PRAD cohort (*n* = 499). Although weaker than the MYC-selected cohorts, negative associations were still observed (Fig. 3D-E).

### MYC negatively regulates MEIS1 expression

Recently, Bhanvadia *et al*., postulated that higher *MEIS1* expression conferred a less aggressive PCa phenotype [19]. Based on these results, we hypothesized that repression of *MEIS1* expression by MYC contributes to MYC-driven PCa. Using LNCaP cells with a non-targeting hairpin as MYC-high and LNCaP cells with three different *MYC*-targeting hairpins as MYC-low (see Supplementary Fig. 1), we performed ChIP-seq against MYC to generate genome-wide site maps and ascertain chromatin occupancy at *MEIS1*. In these cells, we had observed increased overall transcriptional output (see Fig. 1D), and we controlled for this phenomenon using dm3 chromatin spike-in controls. We observed that proportional MYC recruitment to MEIS1 was increased relative to global binding in each of the three *MYC* shRNA lines relative to control (Fig. 4A). Motif enrichment analysis at MYC-ChIP peaks showed modest decreases at canonical MYC binding sites upon *MYC* knockdown (Fig. 4B). We then sought to determine if the observed increase in chromatin occupancy translated to altered MEIS1 transcription by qRT-PCR. Knockdown of *MYC* resulted in increased abundance of the *MEIS1* transcript, consistent with the observed increase in MYC recruitment at *MEIS1* (Fig. 4C). Together, these results support the role of MYC in the negative regulation of *MEIS1* in primary PCa.

**Figure 4.**
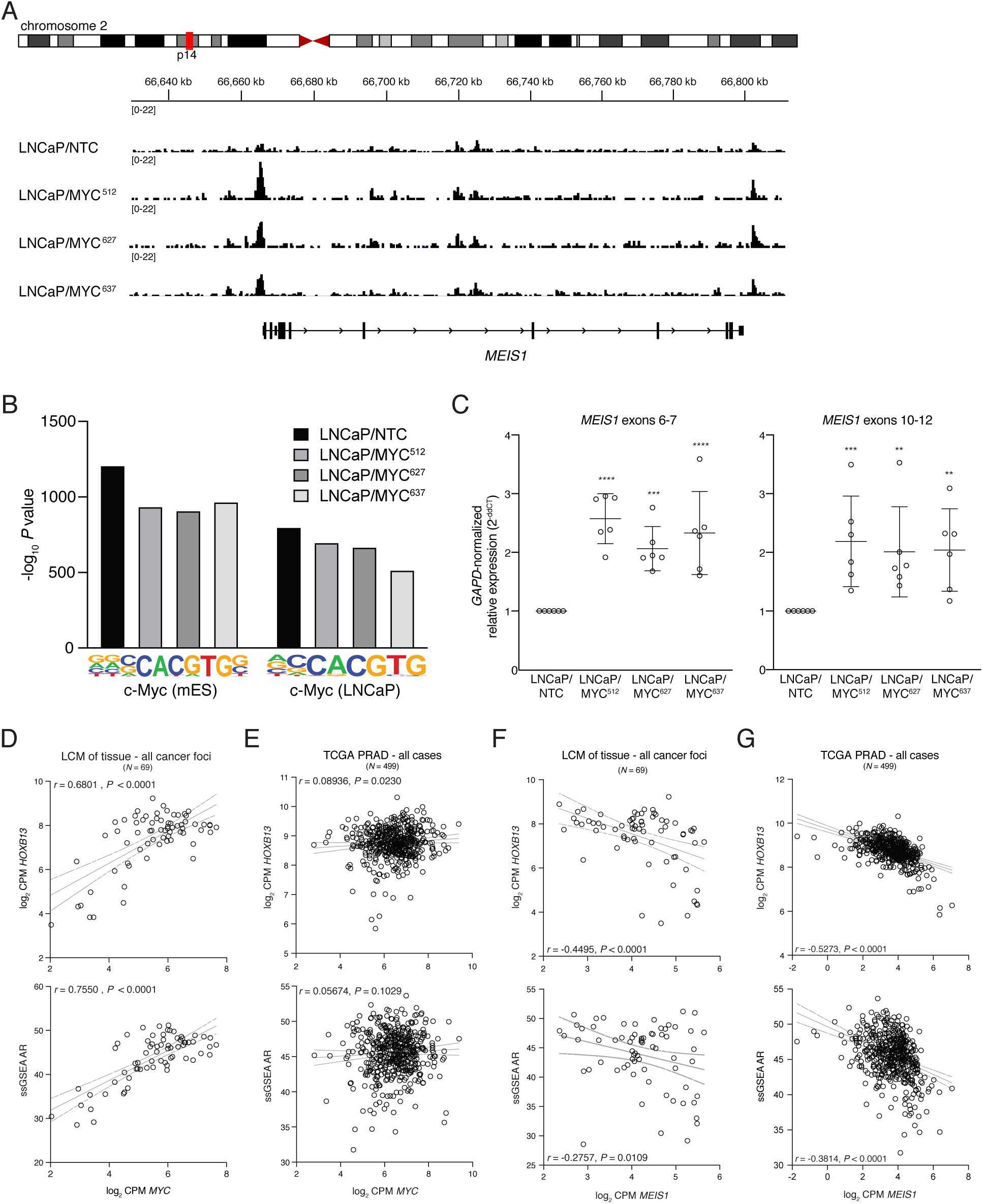
Decreased MYC expression increased MYC occupancy at MEIS1 and MEIS1 expression. A. IGV depiction of MYC binding events in LNCaP *MYC* knockdown cell lines, showing increased MYC occupancy at the *MEIS1* locus in cells harboring *MYC* knockdown (*P* = 0.00513, *Q* = 0.0425, average fold-change: −1.17). B. The confidence of identifying known MYC binding motifs as identified by HOMER are shown for the LNCaP *MYC* knockdown cell lines, measured by −log_10_ *P* value for enrichment. *Q* values (adjusting for false discovery) for these enrichments are zero. C. Quantitative reverse-transcription PCR of *GAPD*-normalized *MEIS1* transcript in LNCaP MYC knockdown cell lines relative to control, measuring across the splice boundary of exons 6-7 (left) or exons 10-12 (right). Bars and whiskers represent the mean ± standard deviation of six independent experiments conducted in triplicate, plotted individually as open circles (**, *P* < 0.01; ***, *P* < 0.001; ****, *P* < 0.0001 by Student’s *t* test). D-E. Correlation of the log_2_ CPM expression level for *MYC* with the log_2_ CPM expression level for *HOXB13* (top) and the 267-gene ssGSEA AR activity signature score (bottom) in the entire LCM cohort (*N* = 69, D) and the entire TCGA PRAD cohort (*N* = 499, E). The gray dotted lines represent the 95% confidence intervals for each correlation. F-G. Correlation of the log_2_ CPM expression level for *MEIS1* with the log_2_ CPM expression level for *HOXB13* (top) and the 267-gene ssGSEA AR activity signature score (bottom) in the entire LCM cohort (*N* = 69, F) and the entire TCGA PRAD cohort (*N* = 499, G). The gray dotted lines represent the 95% confidence intervals for each correlation.

In PCa, MEIS1 functions to direct transcriptional specificity and activity of HOXB13 and act as a negative regulator of AR [20, 45]. Therefore, we next determined whether *HOXB13* expression or AR activity were altered in the context of MYC activity. Indeed, there was a significant positive correlation between *MYC* and *HOXB13* mRNA levels in both the entire LCM tissue cohort (Fig. 4D, *top*) and entire TCGA PRAD cohort (Fig. 4E, top). We also observed a positive correlation between *MYC* mRNA levels and the ssGSEA scores for AR activity [46] in both cohorts, although statistical significance (*P* < 0.05) was reached solely in the LCM cohort (Fig. 4D-E; bottom). Moreover, there was a significant negative correlation between *MEIS1* and *HOXB13* expression in both cohorts (Fig. 4F-G, top) which was also observed between *MEIS1* expression and AR activity (Fig. 4H-I). Taken together, our data suggest that in MYC-high tumors, PCa development is mediated by increased AR activity and *HOXB13* expression resulting from MEIS1 down-regulation.

## DISCUSSION

In many cancer types, the role of amplified MYC in mediating tumorigenesis has been linked to genes involved in ribosomal biogenesis, universally upregulated transcription, proliferation, and reprogramming cells to a pluripotent state [47]. A subset of advanced prostate cancers also harbor amplified MYC, but it is distinct from the upregulated MYC that is a hallmark of many localized prostate cancers [3, 14–16]. In the current study, we used transcriptome profiling to assess subpopulations of prostate tumors based on differential MYC protein expression and MYC activity, and we similarly compared differentially expressed genes and pathways within the larger prostate TCGA cohort based on MYC activity. Finding that increased MYC activity was inversely proportional to overall transcription, we focused on downregulated pathways, identifying a negative correlation between MYC activity and *MEIS1* expression, with MYC directly involved in the negative regulation of *MEIS1* as demonstrated by knockdown and chromatin immunoprecipitation analyses. The inverse association between *MEIS1* expression and AR extended further to *HOXB13* expression, indicating that in a subset of primary PCa, decreased expression of *MEIS1* may be necessary for AR and HOXB13 to drive tumor development and progression.

From its discovery as an oncogene to the subsequent challenges associated with targeting MYC pharmacologically, efforts have shifted in identifying targetable MYC effector genes or other targetable co-factors that are necessary for MYC activity [48, 49]. Efforts to dissect functions of MYC have frequently relied on cancer models in which *MYC* levels rise up to 20-fold, in contrast to 1-2 fold physiological elevation of *MYC* expression in prostate cancer [2, 6, 50]. Not surprisingly, the phenotypes associated with *MYC* up-regulation differ. For example, *MYC* expression in the activation of lymphocytes [36] or in Burkitt’s lymphoma [37] is associated with a universal amplification of transcription, while we observed that in LNCaP prostate cancer cells with up-regulated *MYC*, knocking down *MYC* by less than 50% with shRNA consistently increased the total amount of RNA produced per cell.

The differences in MYC function in prostate cancer extend to the long-standing relationship between MYC and proliferation [47]. Here, we report a series of prostate cancer tissues, serially-sectioned and stained with anti-MYC and anti-Ki67 antibodies, that show a weak proportional relationship. While up to 50% of luminal prostate cancer cells were positive for MYC expression, proliferation measured by Ki67 affected less than 5% of cells. We show a similar relationship in the prostate TCGA cohort and prostate cancer cell lines, suggesting that while proliferating cells may harbor MYC activity, MYC alone is not sufficient for proliferation in prostate cancer. This is generally in agreement with findings that MYC expression alone is not reflective of an increased proliferative fraction of prostate cancer cells [5].

A key finding from our study is the negative relationship between MYC activity and *MEIS1* expression. In the context of MYC’s role as a universal transcriptional amplifier, a target gene of MYC-mediated repression could simply considered a technical anomaly reflecting unequal numbers of cells used for analysis [36]. However, we show that transcriptional signatures of MYC activity and *MEIS1* are inversely correlated in two large independent cohorts, which is largely consistent with the finding that tumors and prostate cancer cells with increased *MEIS1* expression show decreased enrichment of MYC target gene sets by GSEA [19]. Importantly, we further demonstrate here with ChIP-seq that the effects on *MEIS1* expression are due in part to increased MYC occupancy at the *MEIS1* locus, such that lower levels of *MYC* resulted in repositioning of MYC at specific sites. This finding is in disagreement with the model that increased *MYC* expression may have strengthened its activity at a MYC-recruited repressor [36].

Previously, Bhanvadia *et al*. [19] reported that tumors with increased *MEIS1* are potentially less aggressive, based on studies of LAPC-4 prostate cancer cells expressing shRNA against *MEIS1* and that tumors with lower levels of *MEIS1* were at greater risk of biochemical recurrence. HOXB13, a homeodomain transcription factor, has been shown to regulate AR activity while shRNA against *HOXB13* in LAPC4 cells inhibits their growth [18]. Based on these prior findings, as HOXB13 physically interacts with MEIS1 [18], tumors expressing less *MEIS1* would be expected to display greater *HOXB13* expression and AR activity. Indeed, we show an inverse relationship between *MEIS1* expression and AR activity in two independent cohorts, which is consistent with previous observations of a positive correlation between *MYC* expression and AR activity [7].

In summary, our analysis of MYC-expressing prostate tumors demonstrates an inverse relationship with *MEIS1* expression, which in turn is negatively correlated with *HOXB13* expression and AR activity. Mechanistically, our data demonstrate that *MEIS1* is a directly repressed target of MYC, and via effects on *HOXB13* link MYC activity to AR activity. The potential clinical significance of the inverse *MYC*/*MEIS1* relationship warrants further investigation as AR-directed therapies are introduced earlier in the clinical course of disease, and *MEIS1* levels may indicate potential sensitivity to treatment.

## AUTHOR CONTRIBUTIONS

Design of the study: N.C.W., B.E.G., B.J.C., A.G.S.; Tissue collection and pathology: H.Y.; Immunohistochemistry and microdissection: N.C.W., S.Y.T., S.W., R.L., N.V.C., R.A., E.D.W.; Data collection: N.C.W., N.T.T., S.T.H., R.L., N.V.C., R.A., Library preparation and NGS: N.C.W., S.Y.T.; Data analysis: N.C.W., S.Y.T., N.T.T., B.E.G., B.J.C., H.Y., A.G.S.; Manuscript preparation: N.C.W., A.G.S.

## DATA AVAILABILITY

Human tissue RNA-seq data has been deposited into the Database of Genotypes and Phenotypes (https://dbgap.ncbi.nlm.nih.gov/), accession ID phs001813.v1.p1. Human tissue gene expression data has been deposited into the Gene Expression Omnibus (https://www.ncbi.nlm.nih.gov/geo/), accession ID GSE130046. ChIP-seq data has been deposited into the Sequence Read Archive (https://www.ncbi.nlm.nih.gov/sra), accession ID SRP218384 and GEO, accession ID GSE135942.

## ACKNOWLEDGMENTS

The authors gratefully acknowledge the patients and the families of patients who contributed to this study. RNA-seq and ChIP-seq was performed at the CCR Illumina Sequencing Facility. The authors acknowledge technical assistance from Carla Calagua. Portions of this research utilized the computational resources of the NIH HPC Biowulf cluster.

