## Supplementary figures and images for "MEIS1 down-regulation by MYC mediates prostate cancer development through elevated HOXB13 expression and AR activity"

### Supplementary Figure 1

#### Supplementary Figure 1

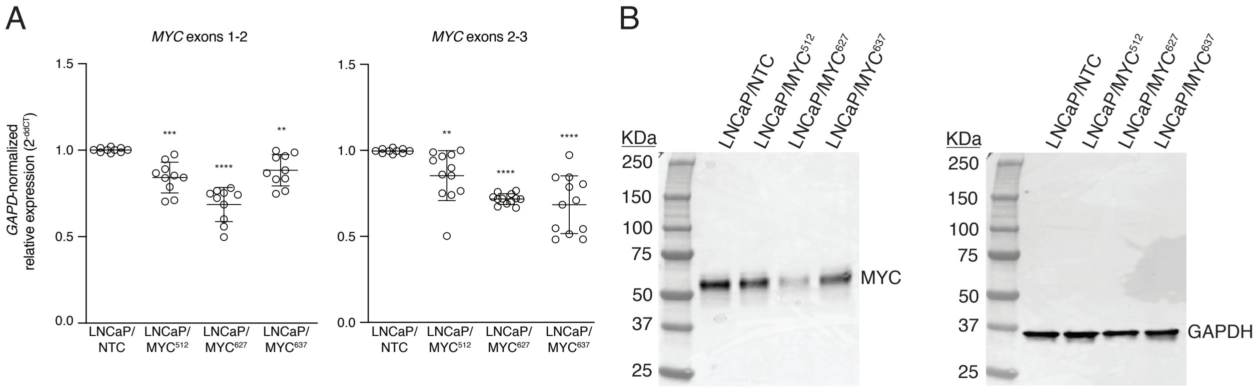

### Supplementary Figure 2

Supplementary Figure 2

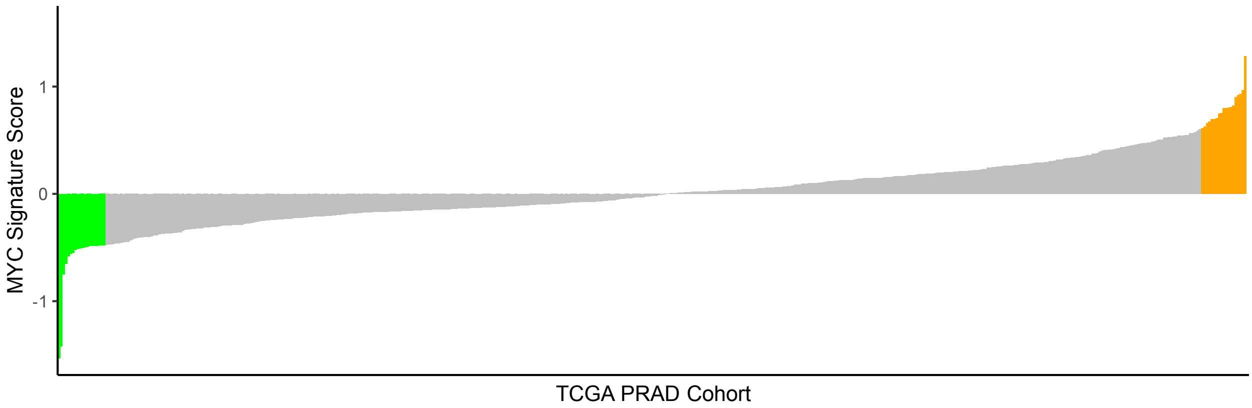
