## Supplementary Information for "MEIS1 down-regulation by MYC mediates prostate cancer development through elevated HOXB13 expression and AR activity"

- **SUPPLEMENTARY TABLES**
- **SUPPLEMENTARY FIGURE LEGENDS**

**SUPPLEMENTARY TABLES**

| **Gene** | **Primer/Probe** | **Sequence** |
| --- | --- | --- |
| *MYC* (exon 1-2) | |  |
|  | Primer 1 | 5′-CAGTAGAAATACGGCTGCAC-3′ |
|  | Primer 2 | 5′-TTCGGGTAGTGGAAAACCAG-3′ |
|  | Probe | 5′-6-FAM-CCGCGACGATGCCCCTCAA-TAM-3′ |
| *MYC* (exon 2-3) | |  |
|  | Primer 1 | 5′-TCTTCCTCATCTTCTTGTTCCTC-3′ |
|  | Primer 2 | 5′-TCCTCGGATTCTCTGCTCTC-3′ |
|  | Probe | 5′-6-FAM-TGGGCGGTGTCTCCTCATGG-TAM-3′ |
| *MEIS1* (exon 6-7) | |  |
|  | Primer 1 | 5′-TGGTGATAGACGATAGAGAAGGA-3′ |
|  | Primer 2 | 5′-ATGCCGTGTCATCATGATCTC-3′ |
|  | Probe | 5′-6-FAM-CCAAGAGGGCTGGTCAGTTAGATTTGC-TAM-3′ |
| *MEIS1* (exon 10-12) | |  |
|  | Primer 1 | 5′-AGAATAGTGCAGCCCATGATAG-3′ |
|  | Primer 2 | 5′-TTCCACTCATAGGTCCTGGT-3′ |
|  | Probe | 5′-6-FAM-CCTTGACTTACTGCTCGGTTGGACT-TAM-3′ |

**Supplementary Table 1**

**Supplementary Table 1.** Sequences of primer-probe sets used for qPCR.

**Supplementary Table 2**

See Microsoft Excel file.

**Supplementary Table 2.** Differentially expressed genes from laser capture microdissected tissues comparing MYC-high versus MYC-low protein expression by immunohistochemistry are ranked by average log_2_ fold-change values, shown with a -log_10_ false-discovery rate that was calculated using the Benjamini-Hochberg correction of a quasi-likelihood framework F test with edgeR.

**Supplementary Table 3**

See Microsoft Excel file.

**Supplementary Table 3.** Differentially expressed genes from the prostate TCGA comparing MYC-high versus MYC-low activity by gene expression signature scoring are ranked by average log_2_ fold-change values, shown with a -log_10_ false-discovery rate that was calculated using the Benjamini-Hochberg correction of a quasi-likelihood framework F test with edgeR.

**Supplementary Table 4**

See Microsoft Excel file.

**Supplementary Table 4.** Overlap of differentially-expressed genes from Supplementary Tables 2 and 3. Statistical significance is based on a false-discovery rate cutoff of 0.1 for tissue and 0.05 for TCGA.

**Supplementary Table 5**

See Microsoft Excel file.

**Supplementary Table 5.** Overlap of significantly enriched mSigDB gene sets based on gene set enrichment analysis of MYC high versus MYC low tissue and TCGA log_2_ fold-change gene expression values, containing at least one differentially expressed gene from Supplementary Tables 2 and 3. Data shown reflect an enrichment *Q* value cut-off of 0.05.

**SUPPLEMENTARY FIGURE LEGENDS**

*Supplementary Figure 1. Knockdown of MYC in LNCaP cells with shRNA.*

A. Quantitative reverse-transcription PCR of *GAPD*-normalized *MYC* transcript in LNCaP *MYC* knockdown cell lines relative to control, measuring across the splice boundary of exons 1-2 (left) or exons 2-3 (right). Bars and whiskers represent the mean ± standard deviation of ten independent experiments conducted in triplicate, plotted individually as open circles (**, *P* < 0.01; ***, *P* < 0.001; ****, *P* < 0.0001 by Student’s *t* test). B. Representative uncropped immunoblots of MYC knockdown LNCaP cell lysates with anti-MYC and anti-GAPDH. Protein molecular weight is given by marker sizes on the left.

*Supplementary Figure 2. Depiction of TCGA cases with high or low MYC activity.*
All prostate TCGA cases were evaluated with an average median absolute deviation-modified *z*-score to determine individual sample MYC activity scores from mRNA expression data. Cases are ranked low to high, with the lowest scoring 20 samples in green and the highest scoring 20 samples in orange.
